# A Prototype for Modular Cell Engineering

**DOI:** 10.1101/170910

**Authors:** Brandon Wilbanks, Donovan S. Layton, Sergio Garcia, Cong T. Trinh

## Abstract

When aiming to produce a target chemical at high yield, titer, and productivity, various combinations of genetic parts available to build the target pathway can generate a large number of strains for characterization. This engineering approach will become increasingly laborious and expensive when seeking to develop desirable strains for optimal production of a large space of biochemicals due to extensive screening. Our recent theoretical development of modular cell (MODCELL) design principles can offer a promising solution for rapid generation of optimal strains by coupling a modular cell and exchangeable production modules in a plug-and-play fashion. In this study, we experimentally validated some designed properties of MODCELL by demonstrating: i) a modular (chassis) cell is required to couple with a production module, a heterologous ethanol pathway, as a testbed, ii) degree of coupling between the modular cell and production modules can be modulated to enhance growth and product synthesis, iii) a modular cell can be used as a host to select an optimal pyruvate decarboxylase (PDC) of the ethanol production module and to help identify a hypothetical PDC protein, and iv) adaptive laboratory evolution based on growth selection of the modular cell can enhance growth and product synthesis rates. We envision that the MODCELL design provides a powerful prototype for modular cell engineering to rapidly create optimal strains for synthesis of a large space of biochemicals.

## Introduction

Cellular metabolisms are diverse and complex, encompassing a substantial space of chemicals ^*1, 2*^. Harnessing these cellular metabolisms for biocatalysis provides a promising path to industrialization of biology that can potentially synthesize these chemicals from renewable feedstocks and organic wastes ^*3-5*^. To achieve this, it is necessary to rewire cellular metabolisms to achieve production efficiency in a rapid and controllable fashion.

Metabolic engineering and synthetic biology are shaping industrialization of biology. A variety of model-guided tools have been developed to enable rational strain engineering that guide desirable genetic knock-outs, knock-ins, and up/down expression systems for redirecting metabolic fluxes to desirable engineered pathways, with applications ranging from production of industrially-relevant bulk chemicals to specialty products and drugs ^*6-11*^. Synergistically, synthetic biology has offered a wide range of genetic tools to engineer promoters ^*12-14*^, ribosome binding sites ^*15*^, terminators ^*16, 17*^, plasmid copy numbers ^*18, 19*^, regulatory and sensory elements ^*20-22*^, and genetic circuits ^*23, 24*^ for controlling metabolic fluxes. With advancement in DNA sequencing, gene synthesis, and pathway assembly, strain variants can be rapidly built and subsequently tested for efficient chemical production ^*22, 25-28*^. The current limitation, however, is to screen for a large space of strain variants through multiple design-build-test cycles of strain optimization to achieve a desirable production phenotype ^*5*^. When expanding to produce a large space of desirable chemicals, the current strain engineering approach will become increasingly laborious and expensive.

Modular design offers the most efficient route for rapid and systematic production that has been applied in most aspects of our modern society, from constructing houses and buildings to transportation systems, industrial factories, and intricate communication networks. Remarkably, biological systems also follow modular design principles ^*29-32*^, and exploiting these principles can potentially facilitate rapid and systematic generation of optimal strains to produce a large space of biochemicals ^*1, 33*^, a promising path toward industrialization of biology. Recent advances in metabolic engineering have significantly progressed to enable modular design of synthetic pathways for combinational biosynthesis of chemicals and fuels ^*3, 34-39*^. However, prototypes for modular cell engineering are still underdeveloped for generating the desirable modular (chassis) cell that is most compatible with pathway modules to achieve most desirable production phenotypes with minimal strain engineering efforts.

A theoretical framework for the modular cell (MODCELL) design has recently been developed to enable modular cell engineering ^*40*^. Based on the MODCELL design principles, a production strain is assembled from a modular (chassis) cell and a production module. A modular cell is designed to be auxotrophic, containing core metabolic pathways that are necessary but insufficient to support cell growth and maintenance. To function, the modular cell must couple with an exchangeable production module containing auxiliary pathways that can complement cell growth and enhance production of targeted molecules. The stronger the coupling between a modular cell and exchangeable production modules, the faster the coupled cells grow, consume substrates, and efficiently produce target chemicals ^*1*^. The production modules are required to balance redox, energy, and intracellular metabolites for sustained cellular metabolism during growth and/or stationary phases. Modular cells are designed to rapidly create optimal production strains with minimal strain optimization cycles.

In this study, we experimentally validate some designed properties of MODCELL, a prototype for modular cell engineering. We show that a modular (chassis) cell is required to couple with a production module, using a heterologous ethanol pathway, as a testbed. By varying the strengths of production modules, we illustrate the degrees of coupling between the modular cell and production modules can be modulated to enhance growth and product synthesis. We further demonstrate a modular cell can be used as a host to select an optimal pyruvate decarboxylase (PDC) of the ethanol production module and to help identify a hypothetical PDC protein. Lastly, we illustrate that adaptive laboratory evolution based on growth selection of the modular cell can enhance growth and target product synthesis rates.

## RESULTS AND DISCUSSION

### Characterization of designed properties of a modular cell and production modules

To validate the MODELL design, we first constructed the modular cell TCS095 (DE3), derived from TCS083 ^*41*^, containing 10 genetic modifications including chromosomal disruption of *pta* (encoding phosphate acetyl transferase), *poxB* (encoding pyruvate oxidase), *ldhA* (encoding lactate dehydrogenase), *adhE* (encoding alcohol dehydrogenase), *zwf* (encoding glucose-6-phosphate dehydrogenase), *ndh* (encoding NADH:quinone oxidoreductase II), *frdA* (encoding fumarate reductase), and *sfcA/maeB* (encoding malate enzyme) as well as chromosomal integration of T7 polymerase gene. This modular cell is a prototype of MODCELL1 that is designed to strongly couple with exchange production modules for producing alcohols (ethanol, butanol, isobutanol) and esters (ethyl butyrate, isobutyl butyrate, and butyl butyrate) ^*40*^. Based on the MODCELL design ^*40*^, the MODCELL1 is auxotrophic under anaerobic conditions due to imbalance of redox and precursor metabolites required for cell synthesis. To validate some properties of MODCELL, we focused on design, construction, and characterization of the ethanol production modules (Figure 1).

**Figure 1.**
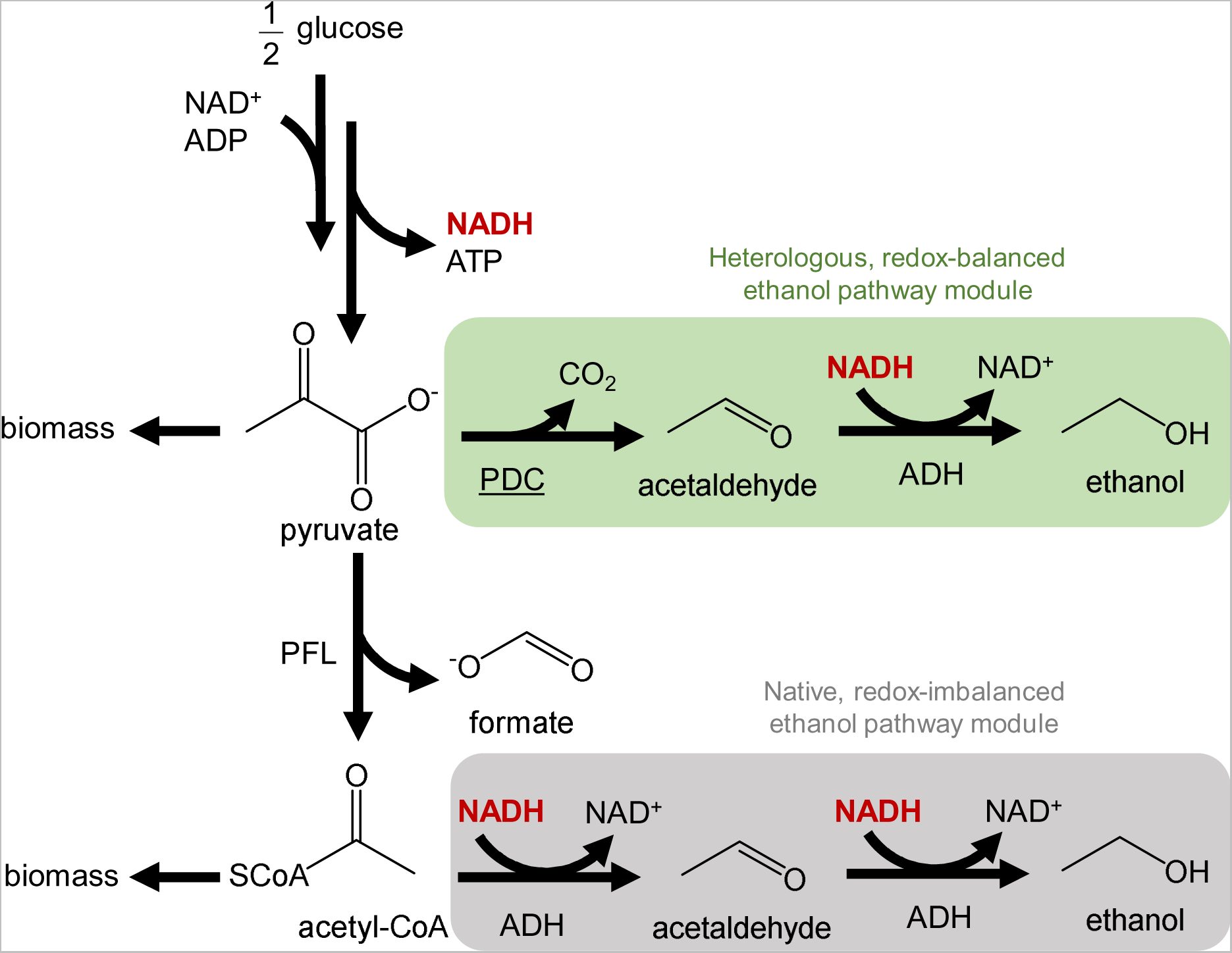
Homoethanol pathways for modular cell growth selection. The pathway highlighted in green is heterologous and redox-balanced that can couple with the modular cell, TCS095 (DE3) or TCS083 (DE3), to enable cell growth under anaerobic conditions. The pathway highlighted in gray is native but redox-unbalanced, which does not enable growth under anaerobic conditions using the modular cell.

We first built the ethanol module, pDL023, a two-operon, two-gene pathway, that is comprised of a pyruvate decarboxylase gene *pdc*_ZM_, derived from *Zymomonas mobilis*, to convert pyruvate to acetaldehyde and an alcohol dehydrogenase gene *adhB*_ZM_ to convert acetaldehyde to ethanol. We also constructed the incomplete ethanol production modules only containing either *pdc*_ZM_ (pCT15) or *adhB*_ZM_ (pCT022). By transforming pCT15, pCT022, and pCT023 into TCS095 (DE3), we generated the coupled cells EcDL107, EcDL108, and ECDL109, respectively (Figure 2A).

**Figure 2.**
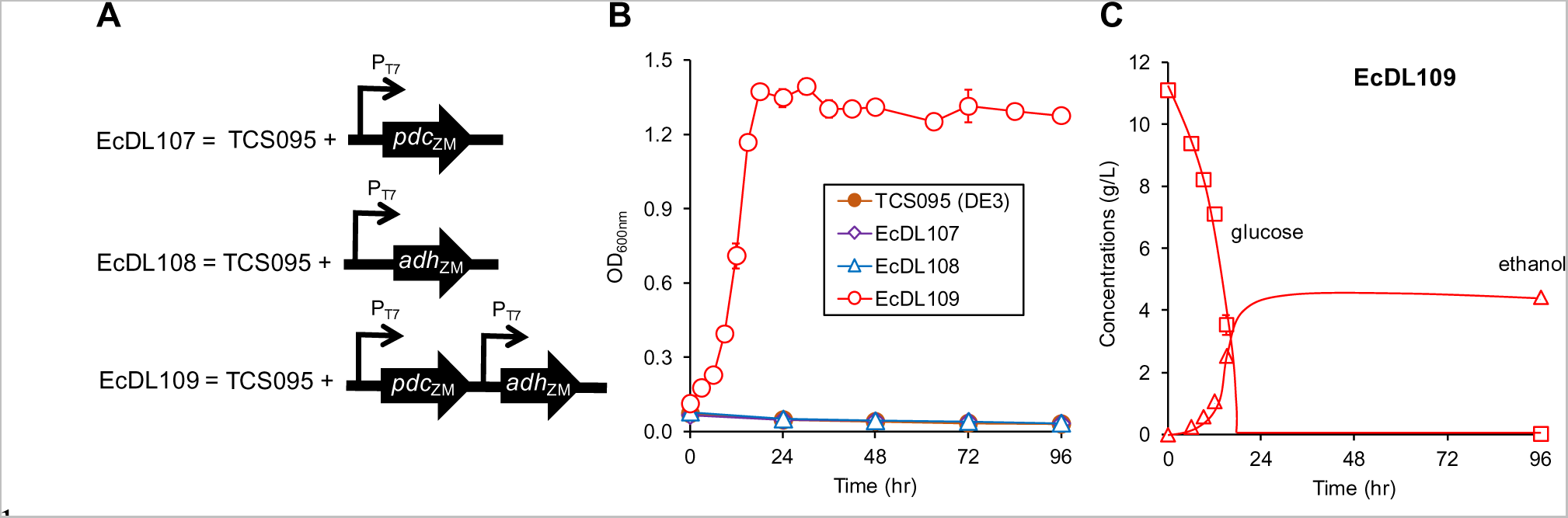
Strong coupling between the modular cell TCS095 (DE3) and ethanol production modules. **(A)** Coupled cells EcDL107, EcDL108, and EcDL109 assembled by the modular cell TCS095 (DE3) and ethanol production modules. **(B)** Cell growth. **(C)** Ethanol production and glucose consumption profiles of EcDL109. The modular cell TCS095 (DE3) and uncoupled cells, EcDL107 and EcDL108, containing incomplete ethanol modules (negative control) while EcDL109 carries a complete two-operon, two-gene ethanol module (test).

Strain characterization shows that the engineered modular cell TCS095 (DE3) indeed could not grow anaerobically while the coupled cell EcDL109 (carrying the complete ethanol production module pDL023) could grow with a specific growth rate of 0.1838 ± 0.0090 (1/h) and reach a maximum optical density (OD, measured at 600nm) of 1.3927 ± 0.0310 (Figure 2B). Furthermore, EcDL109 consumed all glucose within 24 h and mainly produced ethanol with a yield of 0.4507 ± 0.0038 (g ETOH/g GLC) (> 90% of the theoretical limit, 0.51 g ETOH/g GLC) and a specific rate of 0.9131 ± 0.0157 (g ETOH/g DCW/h) (Figure 2C). As a negative control, the uncoupled cells that contained the incomplete ethanol production modules, EcDL107 (carrying pCT15) and EcDL108 (carrying pCT022), could not support cell growth (Figure 2B). Infeasible growth of EcDL107 also implies that AdhB was mainly responsible for conversion of acetaldehyde to ethanol since the native AdhE of the modular cell TCS095 (DE3) was disrupted.

Overall, these results validated the designed property that the modular cell is auxotrophic due to imbalance of redox and precursor metabolites and requires strong coupling with a production module for growth and efficient production of target chemicals. This strategy of strong coupling has been previously shown for production of butanol ^*42, 43*^, isobutanol ^*44*^, succinate ^*45*^, short-chain esters ^*3, 4, 34, 35*^, isopentenol ^*46*^, itaconic acid ^*47*^ from glucose and ethanol from glycerol ^*48*^. Since synthesis and regulation of redox and precursor metabolites are linked within cellular metabolism, any perturbation will affect redox state and precursor requirement for cell growth and maintenance ^*49*^. The redox imbalance resulting in auxotrophy in the modular cell can be readily explained from the simplified metabolic network shown in Figure 1. For the modular cell, the endogenous ethanol pathway is insufficient to maintain redox state because the pathway requires two NADHs per 1/2 glucose while the glycolysis only produces half of the necessary cofactors. Also, strains carrying a deletion of the bifunctional aldehyde/alcohol dehydrogenase AdhE of the ethanol pathway did not enable cellular growth due to underutilization of the NADH cofactor generated from glycolysis. In essence, selection of production modules by the modular cell works as antibiotics but links directly to desirable production phenotypes.

### Modulating degrees of coupling of modular cell and production modules to enhance cell growth and product synthesis

To further demonstrate whether degrees of coupling of the modular cell and production modules can be modulated to enhance cell growth and product formation, we constructed three one-operon, two-gene ethanol modules, with tunable strengths by varying promoters. The strongest module, pCT24, contained the strongest T7 promoter, whereas the weaker modules, pAY1 and pAY3, carried the BBa_J23100 and BBa_J23108 promoters, derived from the iGEM Andersen promoter library ^*50*^, respectively. Among promoters in the library, BBa_J23100 was reported to have the highest strength while BBa_J23108 has 51% activity of BBa_J23100. Due to differences in host strains and growth conditions, we have also independently checked and confirmed the strengths of promoters used in our study: T7 >> BBa_J23100 ≥ BBa_J23108 (Supplementary Figure 1). We constructed three coupled cells, EcDL110, EcDL111, and EcDL112, by transforming the modules pAY3, pAY1, and pCT24 and into the modular cell TCS095 (DE3), respectively (Figure 3A).

**Figure 3.**
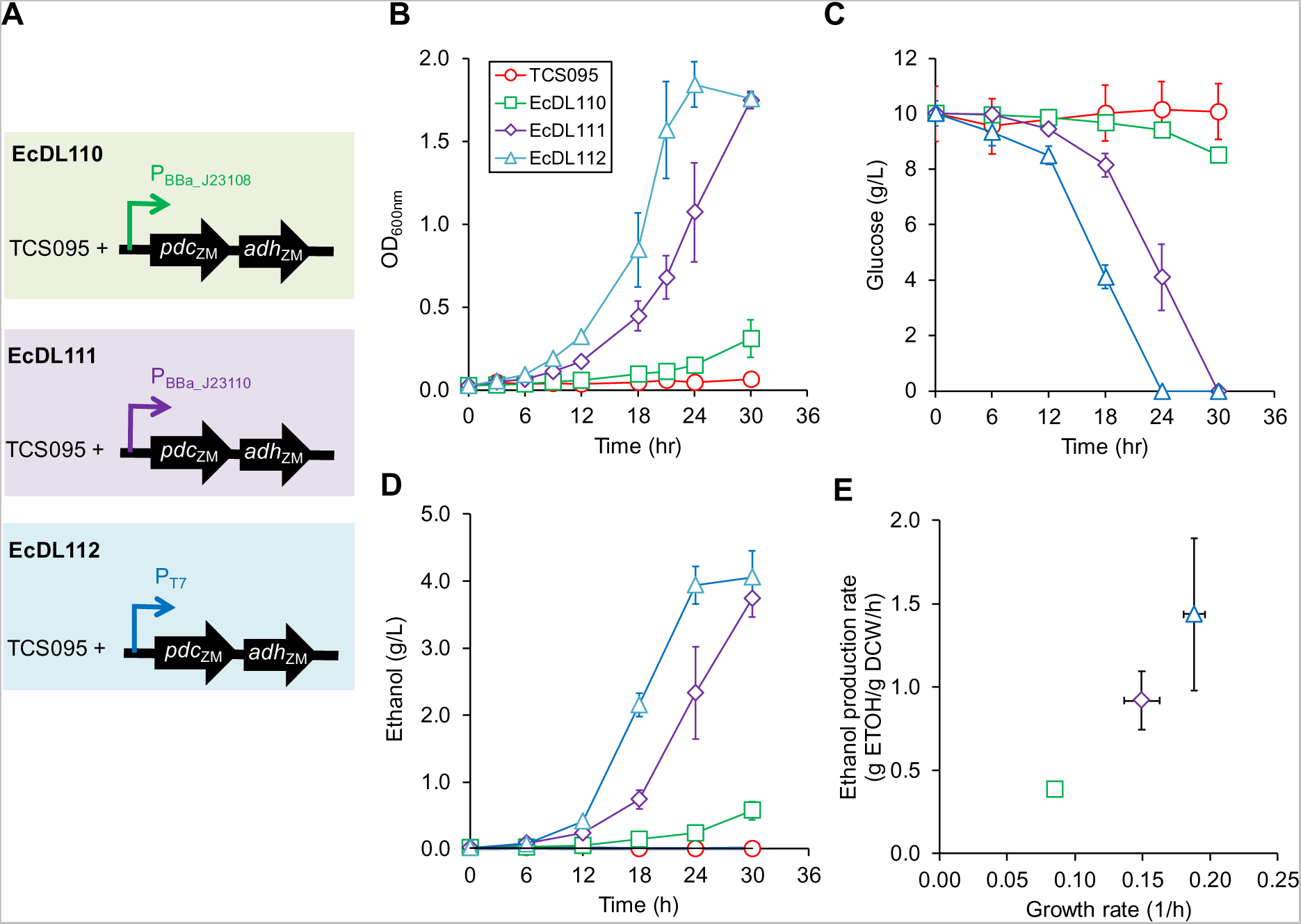
Modulation of degrees of coupling between the modular cell TCS095 (DE3) and ethanol production modules. **(A)** Coupled cells EcDL110, EcDL111, and EcDL112 assembled from the modular cell TCS095 (DE3) and the one-operon, two-gene ethanol production modules, pCT24, pAY1, and pAY3, using promoters of different strengths. **(B)** Cell growth. **(C)** Glucose consumption. **(D)** Ethanol production. **(E)** Correlation of growth and ethanol production rates.

Strain characterization shows that EcDL112, carrying the module pCT24 with the strongest T7 promoter, achieved the highest growth rate of 0.1881 ± 0.0079 (1/h), glucose consumption rate of 3.8216 ± 0.8532 (g GLC/g DCW/h), and ethanol production rate of 1.4350 ± 0.4556 (g ETOH/g DCW/h) (Figure 3B−3D). EcDL111, carrying the module pAY1 with the second strongest promoter, yielded the second highest growth rate of 0.1494 ± 0.0128 (1/h), glucose consumption rate of 2.4799 ± 0.1012 (g GLC/gDCW/h), and ethanol production rate of 0.9176 ± 0.1755 (g ETOH/g DCW/h) (Figure 3B−3D). EcDL110, carrying the module pAY3 with the weakest promoter, obtained the lowest growth rate of 0.0855 ± 0.0007 (1/h), glucose consumption rate of 0.8599 ± 0.2848 (g GLC/g DCW/h), and ethanol production rate of 0.3859 ± 0.0388 (g ETOH/g DCW/h) (Figure 3B−D). Both EcDL112 and EcDL111 consumed all glucose and reached maximum ODs of 1.8433 ± 0.1361 and 1.7500 ± 0.0520, respectively, within 30 h while EcDL110 did not. In addition, we also observed a strong linear correlation (R^2^ = 0.98) between growth and ethanol production rates (Figure 3E).

Taken all together, these results demonstrate that the stronger the coupling between the modular cell and ethanol production modules, the faster the coupled cells grew, consumed glucose, and produced ethanol. Coupling was able to be controlled directly by modifying metabolic fluxes through the pathway of interest. For developing prototypes for modular cell engineering, stronger production modules with balanced fluxes should be enforced to create desirable coupled cells for enhanced production of target chemicals. Production modules with imbalanced fluxes of intermediate steps can significantly affect target product yields as observed for production of isobutanol, ethyl butyrate, isopropyl butyrate, and isobutyl butyrate. ^*3, 4, 34, 35, 44*^

### Enabling modular cells for enzyme selection

#### Selection of PDCs with the modular cell TCS095 (DE3)

Upon confirming the strong coupling between the modular cell and ethanol production module, we next tested whether the modular cell TCS095 (DE3) can be used as a selection host for a target enzyme, i.e., a pyruvate decarboxylase PDC of the ethanol production module. We selected five eukaryotic PDC genes, including *pdc*_Sc1_, *pdc*_Sc5_, and *pdc*_Sc6_ of *S. cerevisiae*, *pdc*_Ppa_ of *P. pastoris*, and putative *pdc*_Yli_ of *Y. lipolytica*, that are divergently different from bacterial PDCs (e.g., *pdc*_ZM_ of *Z. mobilis*). It has been reported that the *in vitro* catalytic efficiencies of PDC_Sc1_, PDC_Sc5_, PDC_Sc6_ and PDC_Ppa_ are relatively similar but significantly lower than that of the bacterial PDC_ZM_ (Supplementary Figure 2) ^*51*^. The activity of putative PDC_Yli_ is widely unknown, which makes it a good candidate to test the selection capability of the modular cell. Should the putative PDC_Yli_ have the predicted function, the coupled cell carrying the PDC_Yli_-dependent ethanol module will grow and produce ethanol; the stronger the activity of PDC_Yli_, the faster the coupled cell will grow and produce ethanol.

We created a library of five two-operon, two-gene ethanol production modules in which we varied the PDC genes and fixed a relatively strong AdhB_ZM_ gene. The complete ethanol production modules, including pDL024, pDL025, pDL026, pDL027, and pDL028, contained *pdc*_Sc1_, *pdc*_Sc5_, *pdc*_Sc6_, *pdc*_Ppa_ and *pdc*_Yli_, respectively, together with the common *adhB_ZM_* gene. By transforming these modules into the modular cell TCS095 (DE3), we constructed the coupled cells EcDL118, EcDL119, EcDL120, EcDL121, and EcDL122, respectively (Figure 4A). For negative controls, we also built the incomplete ethanol modules, including pCT15, pDL017, pDL018, pDL019, pDL020, and pDL021, that contained only *pdc*_Zm_, *pdc*_Sc1_, *pdc*_Sc5_, *pdc*_Sc6_, *pdc*_Ppa_ and *pdc*_Yli_, respectively, without the AdhB_ZM_ gene. The uncoupled cells carrying these incomplete ethanol modules are EcDL107, EcDL113, EcDL114, EcDL115, EcDL116, and EcDL117, respectively (Figure 4A).

**Figure 4.**
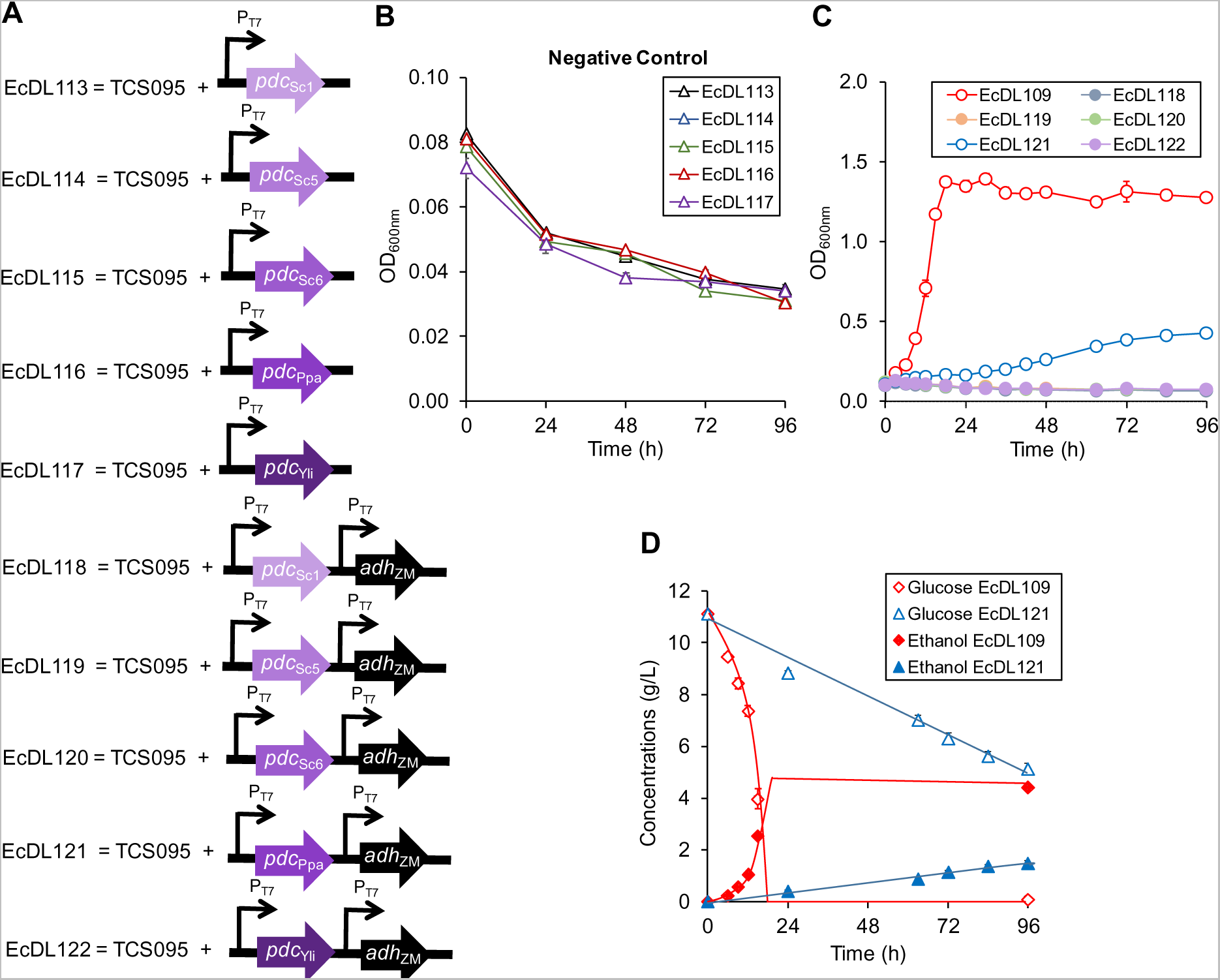
Selection and discovery of pyruvate decarboxylases by the modular cell TCS095 (DE3). **(A)** Uncoupled cells EcDL113-EcDL117 and coupled cells EcDL118-EcDL122 assembled from the modular cell TCS095 (DE3) with incomplete and complete ethanol production modules, respectively. **(B)** Cell growth of the modular cell TCS095 (DE3) and uncoupled cells EcDL113-EcDL117 carrying incomplete ethanol modules only carrying PDCs (negative controls). **(C)** Cell growth of weakly-coupled cells EcDL118-EcDL122 that carry complete ethanol modules with various PDCs. **(D)** Ethanol production and glucose consumption profiles of EcDL109 and EcDL121.

Strain characterization shows that both the negative and positive controls were confirmed. Specifically, the positive control strain EcDL109 grew (Figure 4C) while the negative control strains, including TCS095 (DE3), EcDL107, and EcDL113−EcDL117, could not (Figure 4B). For the test experiments, only the coupled cell EcDL121 containing PDC_Ppa_ grew (Figure 4C). EcDL121 (0.0153 ± 0.0003 1/h) grew much slower than EcDL109 (0.1838 ± 0.0090 1/h) with lower glucose consumption rate (0.5127 ± 0.0097 g GLC/g DCW/h) and ethanol production rate (0.1327 ± 0.0031 g ETOH/g DCW/h) (Figure 4D) mainly because EcDL121 carried a much weaker ethanol production module than EcDL109 (i.e., PDC_ZM_ >> PDC_PPa_). Further, EcDL121 only grew up to a maximum OD of 0.4247 ± 0.0297, ~3.3 fold less than EcDL109, and did not completely consume glucose within 96 h.

Even though some eukaryotic PDC_Sc1_, PDC_Sc5_, and PDC_Sc6_ were previously reported to have higher *in vitro* catalytic efficiencies than PDC_Ppa_ ^*51*^, the coupled cells, EcDL118, EcDL119, and EcDL120, containing these PDCs could not grow. These observed phenotypes could have been caused by non-optimized translation of these eukaryotic PDC genes and/or inefficient *in vivo* enzymatic activities in the heterologous host. As a result, the selection pressure by TCS095 (DE3) might have been too strong for the weak ethanol modules containing these eukaryotic PDCs. The very weak coupling between the modular cell and the modules caused the imbalance of redox and precursor metabolites (e.g., pyruvate, acetyl CoA) and hence inhibited growth of some coupled cells carrying weak eukaryotic PDCs.

#### Selection of PDCs with the modular cell TCS083 (DE3)

To be able to detect the *in vivo* activities of PDC_Sc1_, PDC_Sc5_, PDC_Sc6_, and PDC_Yli_, it is necessary to reduce the strength of selection of TCS095 (DE3). Our strategy was to allow the native, bifunctional acetaldehyde/ethanol dehydrogenase AdhE to relieve redox and/or acetylCoA imbalance that eukaryotic PDCs of the weak ethanol modules alone could not overcome. Based on the MODCELL design, TCS083 (DE3), which is the parent of the modular cell TCS095 (DE3) and still possesses the native AdhE, can function as a modular cell ^*40*^. By coupling TCS083 (DE3) with pDL024 (containing *pdc*_Sc1_/*adhB*_Zm_), pDL025 (*pdc*_Sc5_/*adhB*_Zm_), pDL026 (*pdc*_Sc6_/*adhB*_Zm_), pDL027 (*pdc*_Ppa_/*adhB*_Zm_), and pDL028 (*pdc*_Yli_/*adhB*_Zm_), we constructed the coupled cells EcDL123, EcDL124, EcDL125, EcDL126, and EcDL127, respectively (Figure 5A). The coupled cell EcDL128 carrying pDL023 (*pdc*_Zm_/*adhB*_Zm_) was used as a positive control while the modular cell TCS083 (DE3) was tested as a negative control.

**Figure 5.**
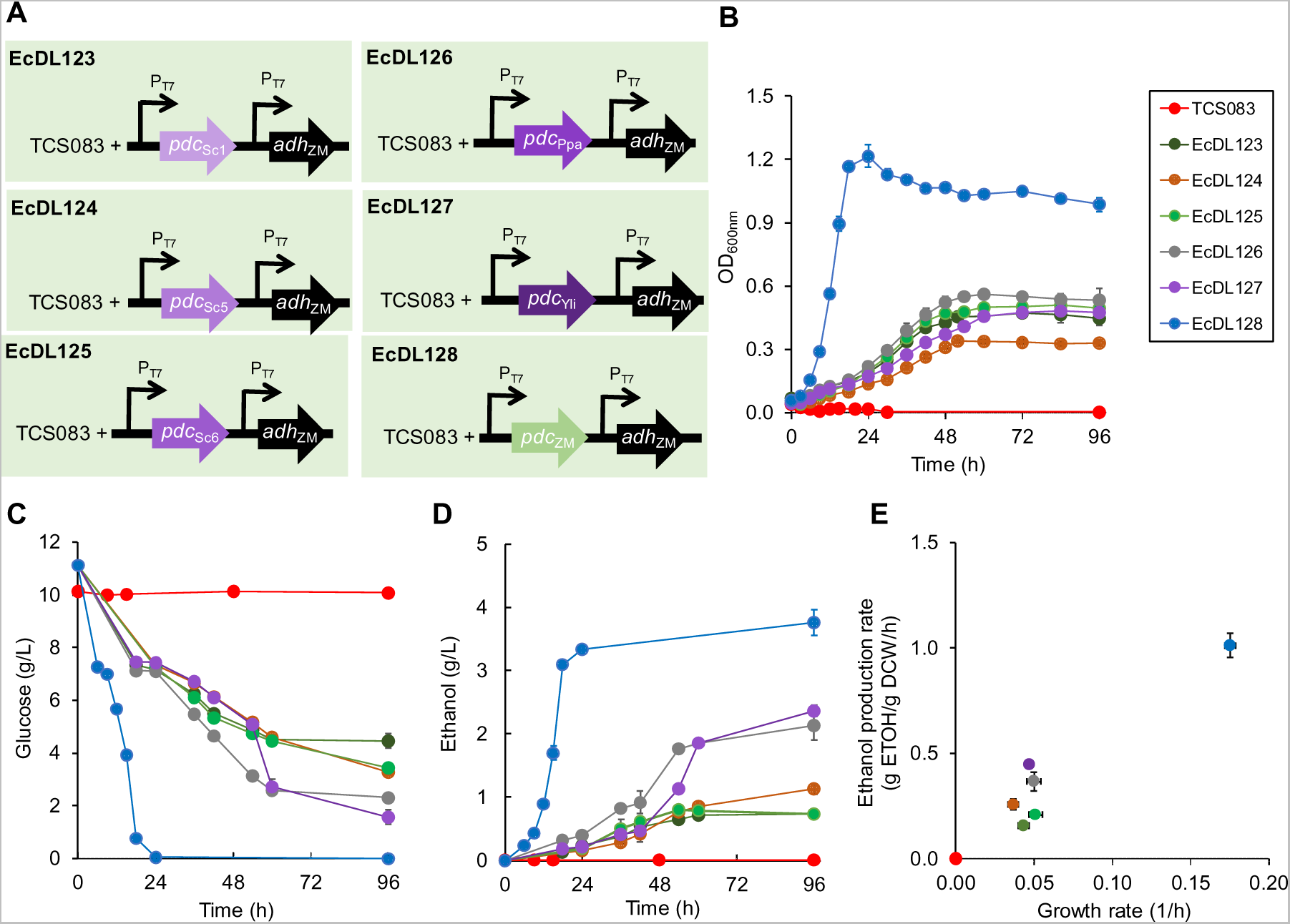
Selection and discovery of pyruvate decarboxylases by the modular cell TCS083 (DE3). **(A)** Coupled cells EcDL123-EcDL128 assembled from the modular cell TCS083 (DE3) with the two-operon, two-gene ethanol modules. **(B)** Cell growth. **(C)** Glucose consumption. **(D)** Ethanol production. **(E)** Correlation of growth and ethanol production rates.

Strain characterization shows that the coupled cell EcDL128, carrying pDL023 (*pdc*_Zm_/*adhB*_Zm_, positive control), could grow while the modular cell TCS083 (DE3) (negative control) alone could not (Figure 5B). As expected, EcDL128, carrying the strongest PDC_Zm_ demonstrated the strongest product coupling among all strains tested with the highest growth rate of 0.1753 ± 0.0034 (1/h), glucose consumption rate of 2.4495 ± 0.0662 (g GLC/g DCW/h), and ethanol production rate of 1.0118 ± 0.0577 g ETOH/g DCW/h (Figure 5B−5E). EcDL128 reached a maximum OD of 1.2167 ± 0.0537 and completely consumed glucose with an ethanol yield of ~ 90% theoretical maximum value (Figure 5). It is interesting to observe that the performance of EcDL128 (derived from TCS083 (DE3)) is very similar to that of EcDL109 (derived from TCS095 (DE3)), which indicates that AdhB alone is sufficiently strong to support the turnover of all necessary acetaldehyde in the designed ethanol production modules.

Using TCS083 (DE3) as the modular cell, all weakly-coupled cells were able to grow. EcDL125, EcDL126, and EcDL127 grew at a similar rate of ~0.05 (1/h), and faster than EcDL123 (0.0434 ± 0.0036 1/h) and EcDL124 (0.0368 ± 0.0033 1/h). As compared to EcDL128, these strains grew ~ 3.4-4.8 fold slower, did not completely consume glucose within 96 h, and reached lower maximum ODs of ~ 0.34-0.56, mainly due to weak ethanol production modules (Figure 5B−5D). Similarly, all TCS083-derived, weakly-coupled cells yielded ethanol production rates of ~2.3-6.4 fold lower than the control EcDL128 (1.0118 ± 0.0577 g ETOH/g DCW/h) (Figure 5E). We also observed a strong correlation (R^2^ = 0.91) between specific growth rates and ethanol production rates of coupled cells carrying ethanol production modules derived from different PDCs. Notably, *Y. lipolytica* PDC is demonstrated for the first time to have *in vivo* activity as observed by ethanol production in EcDL127.

#### Native AdhE in the modular cell TCS083 (DE3) is sufficient to facilitate PDC selection

We further examined whether the native AdhE alone in TCS083 (DE3) is sufficient for the modular cell to select eukaryotic PDCs that are much weaker than PDC_ZM_. We constructed EcDL129, EcDL130, EcDL131, EcDL132, EcDL133, and EcDL134 by introducing pCT15 (containing *pdc*_ZM_), pDL017 (*pdc*_Sc1_), pDL018 (*pdc*_Sc5_), pDL019 (*pdc*_Sc6_), pDL020 (*pdc*_Ppa_), and pDL021 (*pdc*_Yli_) in TCS083 (DE3), respectively (Supplementary Figure 3A). While the modular cell TCS083 (DE3) (negative control) could not grow, all coupled cells could even without using AdhB_ZM_ (Supplementary Figure 3B). This result suggests that the endogenous AdhE was indeed sufficient and required for the ethanol production module to couple with the module cell TCS083.

EcDL129, carrying the strongest PDC_ZM_, grew much faster than the weakly-coupled cells carrying the eukaryotic PDCs by ~2.1-2.7 folds. Interestingly, all weakly-coupled cells EcDL130-EcDL134 exhibited very similar growth, glucose consumption, and ethanol production rates like EcDL123-EcDL127 even though they only used native AdhE without AdhB_ZM_ (Supplementary Figure 3C−3E). In contrast, EcDL129 underperformed EcDL128 when not using AdhB_ZM_. These results suggest that AdhE helped EcDL129-EcDL134 balance redox to support cell growth. The AdhE flux (acetaldehyde + NADH + H^+^ → ethanol + NAD^+^), however, became limiting in EcDL129 but not in EcDL130-134 because PDC_ZM_ of EcDL129 is much stronger than eukaryotic PDCs of EcDL130-EcDL134. Further, the results imply that eukaryotic PDCs were limiting and the bifunctional acetaldehyde/alcohol dehydrogenase AdhE was critical for selection of weak ethanol production modules.

Taken all together, we validate the MODCELL design property that a modular cell can be exploited for enzyme selection and discovery. Like modification of promoter strengths, varying PDCs of the ethanol module provides an alternative method to adjust degrees of coupling between the modular cell and production modules. These couplings can be clearly evidenced by a strong linear correlation (R^2^ = 0.96) between cell growth and ethanol production rates (Supplementary Figure 4A). Manipulating these degrees of coupling also generated some remarkable trends that can help establish a prototype for modular cell engineering.

The coupled cells exhibited optimal growth and ethanol production for a coupling either between a strong modular cell (TCS095 (DE3)) and a strong ethanol module (pDL023 and pCT24) or between a less strong modular cell (TCS083 (DE3)) and a strong ethanol module. In contrast, the coupled cells exhibited significantly slow to no growth and reduced ethanol production phenotypes for a coupling between a strong modular cell and a weak ethanol module (pDL024 − pDL028). Growth and ethanol production were slightly improved for a coupling between a less strong modular cell and a weak ethanol module (EcDL129 − EcDL134). These results underline the importance of balancing push-and-pull carbon and electron fluxes. It is also critical to utilize strong production modules for modular cell engineering to pull carbon and electron fluxes from the core pathways of the modular cell to production modules.

### Adaptive laboratory evolution of the coupled modular cells

#### Evolution of weakly coupled cells resulted in enhanced growth and ethanol production rates

Due to the strong coupling between the modular cell and production modules, we examined whether the adaptive laboratory evolution ^*52, 53*^ based on the growth selection of the modular cell could enhance growth and ethanol production rates of the weakly-coupled cells EcDL130 − EcDL134. We performed the evolution by continuously transferring cultures of EcDL130 − EcDL134 during logarithmic cell growth though serial dilution in two biological replicates for ~ 150 generations (~45 transfers) (Figure 6A). For dilutions 12 (~35 generations), 20 (~60 generations), and 40 (135 generations), we also performed the extensive irreversibility testing by isolating individual colonies of coupled cells and characterizing their performances for cell growth and ethanol production.

**Figure 6.**
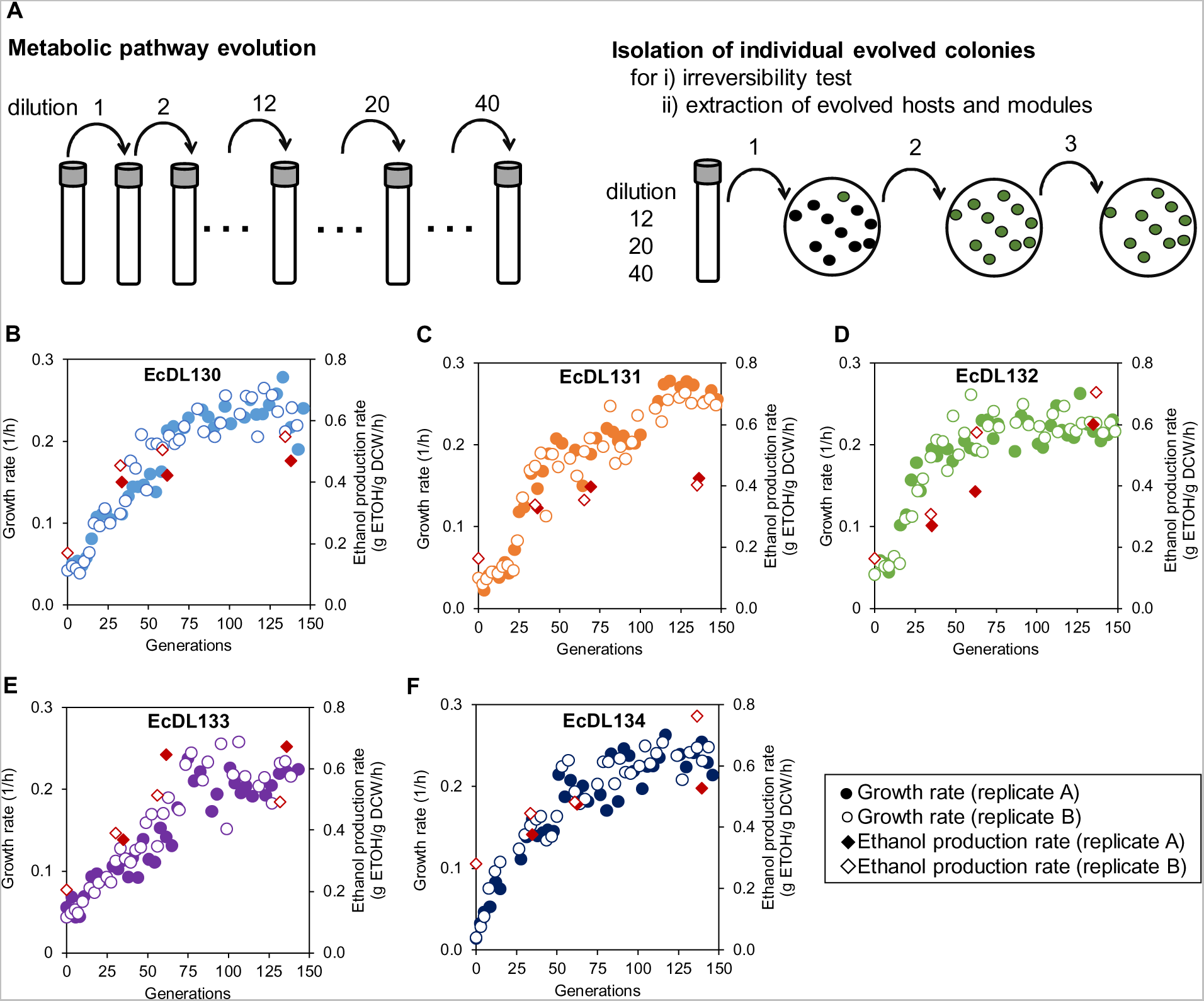
Metabolic pathway evolution enhanced growth and ethanol production rates of weakly-coupled cells. **(A)** Metabolic pathway evolution carried out by the serial culture dilution. Individual evolved cells can be isolated by serial plate spreading. Characterization of growth and ethanol production rates during the metabolic pathway evolution of weakly coupled cells including **(A)** EcDL130, **(B)** EcDL131, **(C)** EcDL132, **(D)** EcDL133, and **(E)** EcDL134 for a period of 150 generations. In panels **B-F**, the ethanol production rates were evaluated from individual isolates from dilutions 12, 20, and 40.

The results show that the evolved EcDL130e − EcDL134e significantly improved growth and ethanol production rates during the adaptive laboratory evolution (Figure 6B−6D). It should be noted that the letter “e” following EcDL130 − EcDL134 signifies that these strains EcDL130e-EcDL134e undergo an adaptive laboratory evolution; and when a number appears after “e”, it represents the number of dilution. For instance, EcDL130e40 is an evolved strain isolated at the dilution 40. For comparison and discussion hereafter, we used the performance of EcDL129 (μ = 0.1021 ± 0.0059 1/h and r_P_ = 0.4819 ± 0.0368 g ETOH/g DCW/h) as a benchmark because it accomplished the highest growth and ethanol production rates of all six PDCs tested before the evolution.

Initially, the weakly-coupled cells EcDL130−EcDL134 grew slowly in a range of 0.04-0.05 1/h. After dilution 12, all cells significantly improved growth by ~2-4 folds, and either reached or surpassed the growth of EcDL129. Specifically, EcDL132e12 grew fastest with a growth rate of ~1.9 folds higher than the benchmark strain EcDL129. After dilution 20, all cells almost doubled the growth rate of the benchmark strain EcDL129 where specific growth rates of EcDL130e20, EcDL131e20, EcDL132e20, EcDL133e20, and EcDL134e20 were 2.0, 1.9, 1.9, 1.6 and 2.2 folds higher than that of EcDL129, respectively. After dilution 20, coupled cells EcDL130e40−EcDL134e40 slightly improved growth where growth rates mostly reached plateau at dilution 40, ~2.0-2.6 folds higher than EcDL129. Single colony isolates at dilutions 12, 20, and 40 were also characterized for irreversibility test, and all matched the observed, enhanced growth phenotypes of evolved cultures (Supplementary Figure 5). Throughout dilution, evolved cells enhanced not only growth but also glucose consumption and ethanol production rates (Supplementary Figure 6). At dilution 40, coupled cells EcDL130e40, EcDL131e40, EcDL132e40, EcDL133e40, and EcDL134e40 reached the ethanol production rates ~3.0, 2.5, 4.0, 2.8, and 2.3 folds higher than their parents, respectively. All coupled cells either matched or slightly surpassed the ethanol production rate of EcDL129. Interestingly, we observed that there exists a weak correlation between growth and ethanol production rates for EcDL130−134 and their evolved derivatives at dilutions 12, 20, and 40 (Supplementary Figure 4B).

#### Host adaption contributed to enhanced performance of coupled cells

To examine whether the host or production modules contributed to enhanced phenotypes of evolved mutants, we isolated and characterized both. To characterize the evolved ethanol production modules, we selected three representative evolved cells EcDL130e40 (with PDC_Sc1_), EcDL133e40 (with PDC_Ppa_), and EcDL134e40 (with PDC_Yli_), extracted their plasmids, transformed them back to the unevolved parent modular cell TCS083 (DE3), and characterized. The results show that these coupled cells did not improve growth rates as compared to their parents (Supplementary Figure 7).

To test the host, we rejected the plasmids from the evolved cells through serial plate transfers. After multiple rounds of strain curation through plate transfers, we were able to successfully isolate TCS083e40 by rejecting the plasmid pDL020 from EcDL133e40 first and hence used it for characterization. We transformed the original modules pCT15, pDL017, pDL018, pDL019, pDL020, and pDL021 into TCS083e40 for characterization. Most coupled cells were able to regain the enhanced growth phenotypes of the evolved mutants (Figure 7A). Interestingly, TCS083e40 pCT15 doubled the growth rate of EcDL129 (TCS083 pCT15) (Figure 7B). In addition to improved growth, we also observed increase in ethanol production rates of the coupled cells derived from the evolved modular cell TCS083e40 and the original ethanol production modules (Figure 7C).

**Figure 7.**
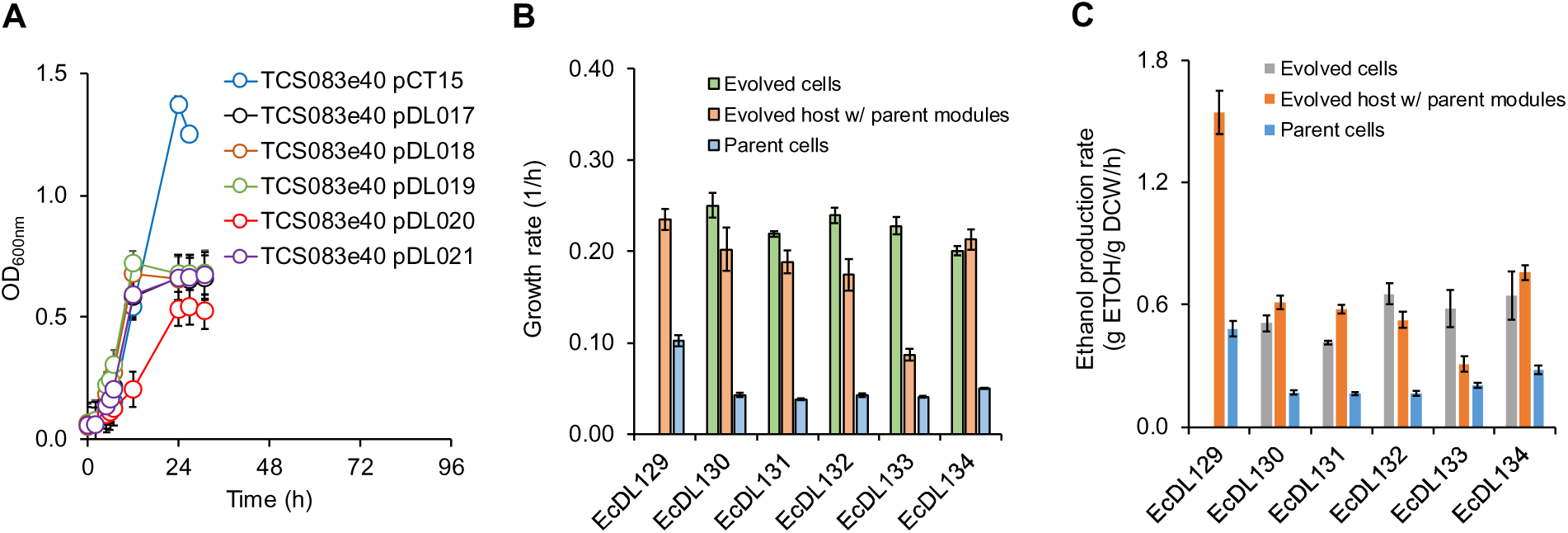
Adapted modular cells contributed to enhanced phenotypes of evolved cells. **(A)** Growth kinetics of the coupled cells assembled from the adapted modular cell carrying the parent ethanol modules including pCT15, pDL017, pDL018, pDL019, pDL020, and pDL021. The evolved modular cell TCS083e30 was isolated from EcDL133e40 at dilution 40 after rejecting pDL020. Comparison of **(B)** growth rate and **(C)** ethanol production rate among the parent coupled cells, evolved cells, and adapted modular cell carrying the parent ethanol production modules.

Taken all together, these results demonstrate that the modular cell can be used a selection host for adaptive laboratory evolution to enhance growth and production synthesis rates of coupled cells. The evolved EcDL130e-EcDL134e might have acquired beneficial mutations on core metabolisms of the modular cell to achieve higher growth and ethanol production rates but probably not on the production modules. Future studies will investigate the beneficial mutations through OMICS analysis and genome resequencing. Our results concur with previous studies of plasmid-bacteria evolution that plasmids tend to resist mutations even undergoing long-term (>500 generations) adaptive laboratory evolution experiments ^*54, 55*^. Based on this observation, to acquire beneficial mutations on the production modules, it might be necessary to generate *in vitro* module variants (e.g., protein mutagenesis ^*56*^) and use the modular cell for selection.

## CONCLUSION

We have developed a prototype for modular cell engineering based on the MODCELL design principles. Using a heterologous ethanol pathway as a testbed, we characterized and validated some designed properties of a modular cell. We demonstrated the auxotrophy of two modular cell designs by coupling them with no module or any incomplete module. By modulating the degrees of coupling with various promoter strengths or activities of PDCs of ethanol production modules, we demonstrated that the strong coupling is critical for enhanced growth and product formation. This strong coupling enabled the modular cell to be used as a host to select an optimal pyruvate decarboxylase (PDC) of the ethanol production module or discover function of the hypothetical PDC protein from *Y. lipolytica*. Using the modular cell platform, adaptive laboratory evolution based on growth selection provides a simple but powerful technique to enhance growth and product rates of a targeted pathway. We envision that MODCELL provides a powerful prototype of modular engineering to rapidly create optimal strains for efficient production of a large space of biochemicals and help minimize the design-build-test cycles of strain engineering.

## METHODS

### Strains

Table 1 lists strains used in this study. *E. coli* TOP10 was used for molecular cloning. TCS095 was constructed from TCS083 ^*41*^ by deleting chromosomal gene *adhE* using P1 transduction ^*57*^. The prophage λDE3 was used to insert a T7 polymerase gene into the specific site of TCS083 or TCS095 by using a commercial kit for strains expressing a T7 promoter (cat#69734-3, Novagen Inc.). TCS083 (λDE3), TCS095 (λDE3), and their derivatives carrying production modules were used for modular cell engineering and characterization (Table 1). All mutants and plasmids were PCR confirmed with the primers used listed in Supplementary Table 1.

**Table 1.**
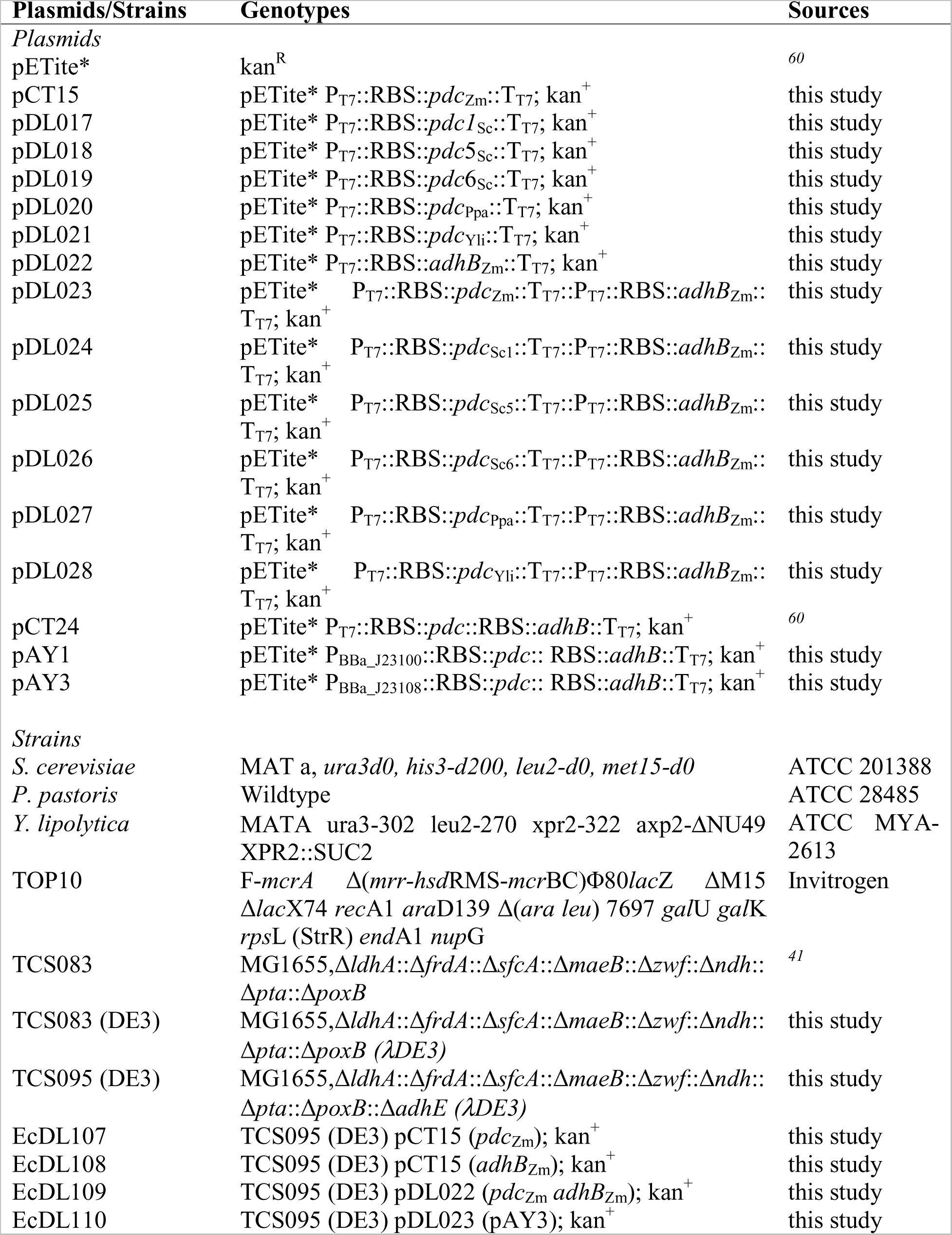
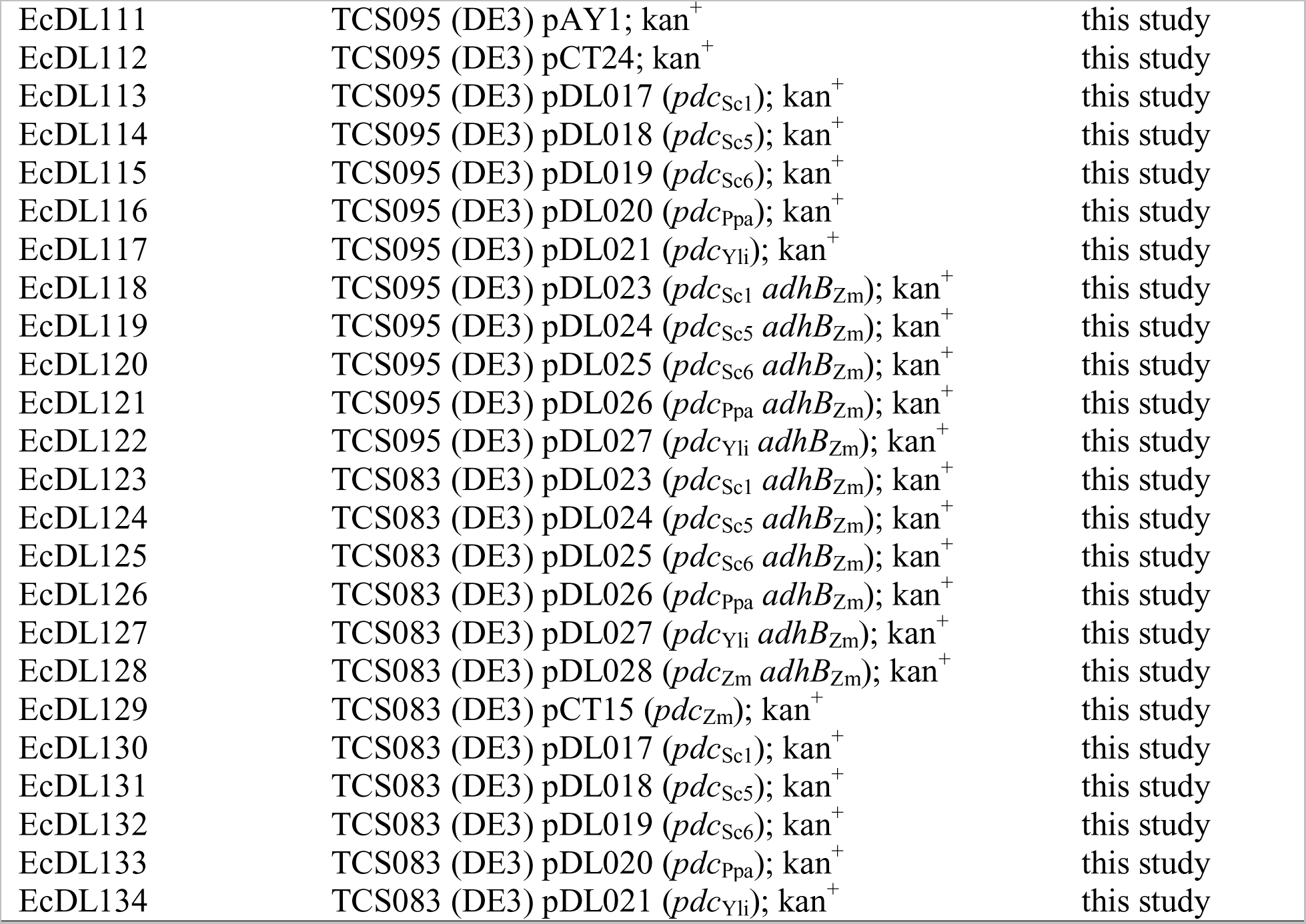
A list of strains and plasmids used in this study.

### Plasmid/pathway construction

#### Construction of PDC modules

The pETite*, a vector backbone ^*34*^, was used to construct PDC modules − containing PDC genes derived from *Z. mobilis*, *S. cerevisiae*, *P. pastoris*, and *Y. lipolytica* under T7 promoters − using the Gibson gene assembly method ^*58*^. To construct the modules pDL017, pDL018, and pDL019, the genes PDC1_Sc_, PDC5_Sc_, and PDC6_Sc_ were amplified from *S. cerevisiae* cDNA using the primers DL_0036/ DL_0037, DL_0038/ DL_0039, and DL_0040/ DL_0041, respectively, and then inserted into the pETite* backbone isolated by using the primers DL_0001/DL_0002. Likewise, the PDC_Ppa_ gene was amplified from *P. pastoris* cDNA using the primers DL_0042/ DL_0043 and inserted into the pETite* backbone, generating the module pDL020. The module pDL021 was constructed by amplifying the PDC_Yli_ gene from *Y. lipolytica* cDNA using the primers DL_0044/ DL_0045 and inserted into the pETite* backbone.

The *Z. mobilis* PDC_ZM_ gene was assembled into the pETite* vector to create the module pCT15 by using the BglBrick gene assembly ^*59*^ of 2 DNA pieces: i) PDC_ZM_ gene amplified using genomic DNA of *Z. mobilis* using the primers P006_f/P006_r and digested with NdeI/BamHI and ii) the vector backbone pETite* doubly digested with NdeI/BamHI.

#### Construction of AdhB modules

The AdhB module pDL022 was generated by amplifying AdhB_Zm_ from pCT24 using primers DL_0046/ DL_0047 and inserted into the pETite* backbone using Gibson assembly.

#### Construction of the PDC and AdhB ethanol modules

The single-operon, PDC/AdhB ethanol module pCT24 was previously constructed ^*60*^. Briefly, pCT24 contains the *Z. mobilis* ethanol pathway encompassing PDC and AdhB genes. The variant PDC/AdhB ethanol modules pAY1 and pAY3 were constructed from pCT24 by swapping the T7 promoter with two weaker constitutive promoters ^*50*^. The module pAY1 contains the BBa_J23100 promoter that was constructed by amplifying pCT24 using the primers AY6.R/AY.7F, digested with BglII, and ligated together. Likewise, the module pAY3 carries the BBa_J23108 promoter that was constructed using the primers AY6.R/AY10.F, digested with BglII and ligated together.

In addition, we have constructed and characterized the two-operon, PDC/AdhB modules: the first operon carrying a PDC gene derived from various species and the second operon containing the adhB_Zm_ gene. The two-operon ethanol modules were constructed using the Gibson assembly method using two parts: i) the PDC containing-plasmids pDL017-pDL021, pCT15 amplified using primers DL_0013/DL_0014 and ii) the adhB operon from pDL022 using primers DL_0015/DL_0016 to generate pDL023-pDL028, respectively.

### Medium and cell culturing

#### Culture media

For molecular cloning, the lysogeny broth (LB) medium, containing 10 g/L yeast extract, 5 g/L tryptone, and 5 g/L NaCl, was used. Antibiotics at working concentrations of 50 μg/mL kanamycin (kan) was used, where applicable, to maintain the selection of desired plasmids. For growth coupling experiments, the M9 (pH~7) medium was used, consisting of 100 mL/L of 10X M9 salts, 1 ml/L of 1 M MgSO_4_, 100 μL/L of 1M CaCl_2_, 1 ml/L of stock thiamine HCl solution (1 g/L), 1 ml/L of stock trace metals solution ^*41*^, and appropriate antibiotics. Unless specified, 10 g/L glucose was used in the M9 medium. The stock 10X M9 salt solution contained 67.8 g/L Na_2_HPO_4_, 30 g/L KH_2_PO_4_, 5 g/L NaCl, and 10 g/L NH_4_Cl.

#### Strain characterization

Strain characterization experiments were performed by growing cells overnight at 37^o^C in 15mL culture tubes containing LB and appropriate antibiotics, then subculturing into fresh M9 medium to adapt the cells to a defined environment. Cells were then grown until exponential phase (OD_600nm_ ~1.0, 1 OD ~ 0.5 g DCW/L). Next, cells (except the modular strain TCS083 DE3) were again subcultured into a nitrogen sparged and pressured tube to create a complete anaerobic environment to an initial OD_600nm_ ~0.10-0.20 at a working volume of 20 mL. The strains were allowed to adapt (at least 2 doublings) overnight to the anaerobic environment and then transferred into pre-warmed 20 mL tubes dispersed of oxygen containing M9 and appropriate antibiotics for characterization with an initial OD_600nm_ of ~0.030.

Cells were grown on a 75^o^ angled platform in a New Brunswick Excella E25 at 37^o^C and 175 rpm. Whole-cells and cell supernatants were collected and stored at −20°C for subsequent metabolite analysis. All experiments were performed with at least three biological replicates.

#### Adaptive laboratory evolution

Strain evolution experiments were prepared and grown in an identical way to the method described above for strain characterization experiments. Samples for metabolite analysis were also taken in a similar manner as described in the strain characterization method. Upon preparation of adapted anaerobic strains, cultures were grown in duplicate from OD_600_ of ~0.050 until exponential phase was reached (OD_600_ of 0.5-1.0). A 1.5 mL sample of each replicate was collected for stock. Each replicate was then diluted to OD_600_ ~ 0.05 and grown again to an OD_600_ of 0.5-1.0, where samples were collected as before. This process was repeated until a consistent maximum growth rate was reached. Evolution was then tested for irreversibility.

When the consistent maximum growth rate was reached, cells were plated on LB plates with antibiotic as needed. A single colony was selected and streaked out on a new plate and repeated 3 times in order to ensure a single cell colony was isolated. Isolated evolved colonies were tested for irreversibility. Cells were first grown and stocked at −80°C before conducting the experiment to allow for complete metabolic interruption and recovery. Irreversibility of the adapted host strain and adapted plasmid were carried out in the same fashion as before in the strain characterization method. Plasmids were extracted from the isolated colonies and transformation into the unevolved parent strain (TCS083 DE3) for plasmid irreversibility test. The evolved plasmid in the unevolved host was also characterized in the same method of the strain coupling studies.

### Analytical methods

#### Cell growth

Cell optical density was measured to quantify cell growth by using a Thermo Scientific Genysys 30 Visible Spectrophotometer with a proper adapter to measure growth kinetics directly.

#### High performance liquid chromatography (HPLC)

Extracellular metabolites were quantified by first filtering cell supernatants though 0.2-μm filter units and then analyzed using the Shimadzu HPLC system equipped with RID and UV-Vis detectors (Shimadzu Inc., Columbia, MD, USA) and Aminex HPX-87H cation exchange column (BioRad Inc., Hercules, CA, USA). Samples were eluded though the column set at 50°C with a flow rate of 0.6 mL/min using the 10 mN H_2_SO_4_ mobile phase ^*44*^.

### Data analysis

***Specific growth rate**.* First-order kinetics was applied to calculate a specific growth rate from kinetic measurement of cell growth as follows:

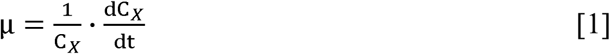

where μ (1/h) is the specific growth rate, C_X_ (g/L) is cell titer, and t (h) is culturing time.

#### Yield

A yield (Y_Si_/Y_Sj_) of a species S_i_ with respect to a species S_j_ (i # j) is determined as follows:

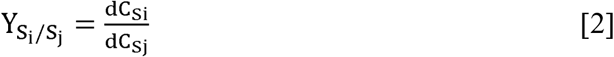

where C_Si_ and C_Sj_ (g/L) are concentrations of S_i_ and S_j_, respectively.

#### Specific production/consumption rate

A specific rate r_Si_ of a species S_i_ is calculated as follows:

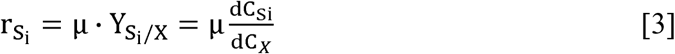

***Growth generation**.* The number of growth generation (n) is determined as follows:

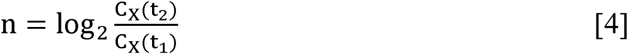

## ACKNOWLEDGEMENTS

We thank Akshitha Yarrabothula, Nirayan Niraula, and Katherine Krouse for helping with molecular cloning and growth study experiments.

## AUTHOR CONTRIBUTIONS

CTT perceived and supervised the study. CTT, DL, BW designed experiments. DL, BW, SG performed the experiments. CTT, DL, BW analyzed the data. CTT wrote the manuscript.

